# Benzimidazoles cause lethality by inhibiting the function of *Caenorhabditis elegans* neuronal beta-tubulin

**DOI:** 10.1101/2022.07.21.500991

**Authors:** Sophia B. Gibson, Elan Ness-Cohn, Erik C. Andersen

## Abstract

Parasitic nematode infections cause an enormous global burden to both human and livestock populations. Resistance to the limited arsenal of anthelmintic drugs used to combat these infections is widespread, including resistance to benzimidazole (BZ) compounds commonly found in livestock parasites. Previous studies using the free-living nematode *Caenorhabditis elegans* to model parasitic nematode resistance have shown that loss-of-function mutations in the beta-tubulin gene *ben-1* confer resistance to BZ drugs. However, the mechanism of resistance and the tissue-specific susceptibility are not well known in any nematode species. To identify in which tissue(s) *ben-1* function underlies BZ susceptibility, transgenic strains that express *ben-1* in different tissues, including hypodermis, muscles, neurons, intestine, and ubiquitous expression were generated. High-throughput fitness assays were performed to measure and compare the quantitative responses to BZ compounds among different transgenic lines. Significant BZ susceptibility was observed in animals expressing *ben-1* in neurons, comparable to expression using the *ben-1* promoter. This result suggests that *ben-1* function in neurons underlies susceptibility to BZ. Subsetting neuronal expression of *ben-1* based on neurotransmitter system further restricted *ben-1* function in cholinergic neurons to cause BZ susceptibility. These results better inform our current understanding of the cellular mode of action of BZ and also suggest additional treatments that might potentiate the effects of BZs.

**Highlights:**

- Expressing wild-type *ben-1* only in neurons restores susceptibility to benzimidazoles
- Expression of *ben-1* in cholinergic neurons restores susceptibility to benzimidazoles
- GABAergic neurons might also play a role in benzimidazole sensitivity
- Broad implications for molecular mechanisms of benzimidazole mode of action

**Graphical Abstract:** 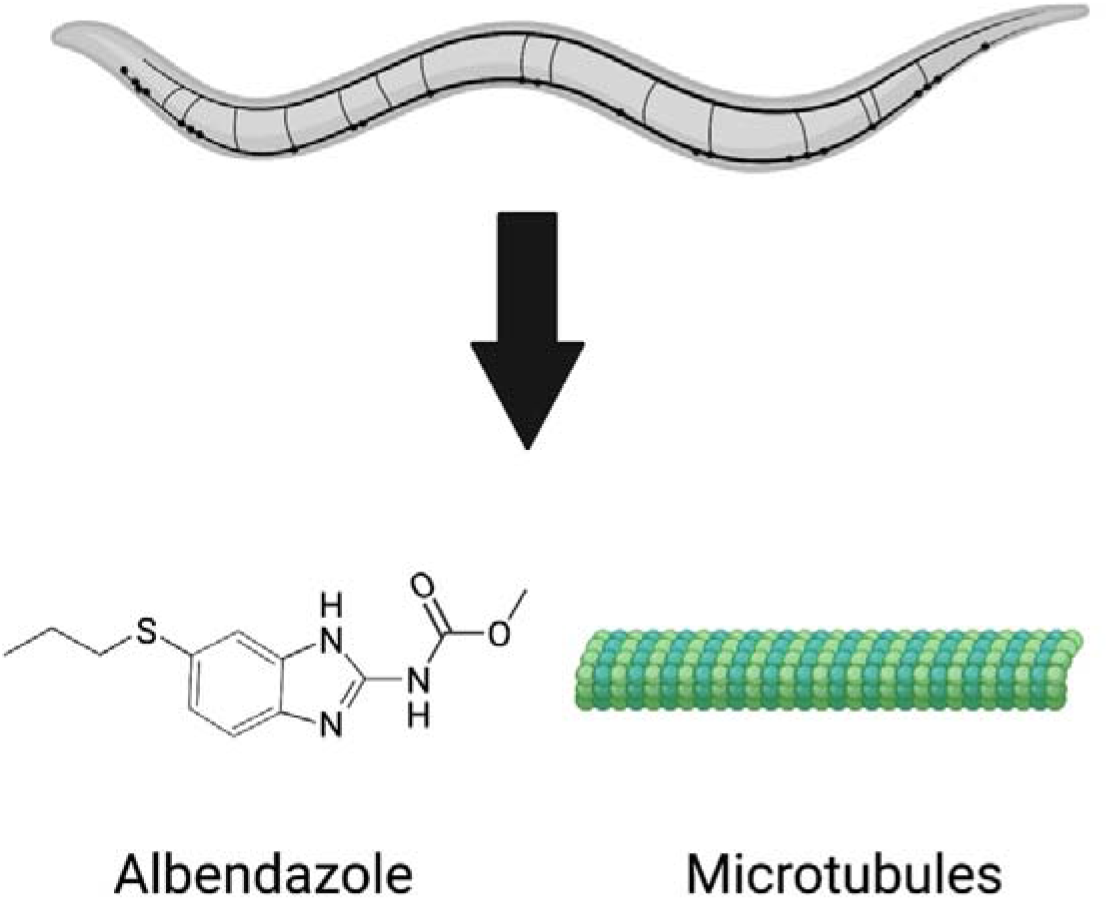

## 1. Introduction

Anthelmintic drugs are crucial to combat parasitic nematode infections, which affect billions of people and livestock populations each year (Hotez et al., 2014; Kaplan & Vidyashankar, 2012). However, only a limited arsenal of drugs are approved, comprising four major classes: benzimidazoles (BZs), nicotinic acetylcholine receptor agonists (nAChRs), macrocyclic lactones (MLs), and amino-acetonitrile derivatives (AADs). BZs have been used extensively for over 50 years (Abongwa et al., 2017; Roos et al., 1995). Because of the intensity of administration of the few anthelmintics available, resistance was documented in *Haemonchus contortus* within a few years of its introduction (Theodorides et al., 1970). Resistance to commonly used BZs continues to be widespread today (Kaplan & Vidyashankar, 2012).

Following the establishment of resistance to BZs, mutations in beta-tubulin genes were first correlated with resistance in the fungus *Aspergillus nidulans*, followed by the free-living nematode *Caenorhabditis elegans* (Driscoll et al., 1989; Hastie & Georgopoulos, 1971; Sheir-Neiss et al., 1978). Loss-of-function mutations in the beta-tubulin gene *ben-1* were identified in strains resistant to BZs (Driscoll et al., 1989). Redundancy among the six beta-tubulin genes in *C. elegans* allows strains with non-functional *ben-1* to still develop normally (Driscoll et al., 1989). Resistance alleles corresponding to point mutations in *ben-1* homologs in parasitic nematode populations continue to be identified (Avramenko et al., 2019; Dilks et al., 2021; Hahnel et al., 2018; Mohammedsalih et al., 2020), and many have been validated to cause resistance using genome editing and highly sensitive *C. elegans* drug response assays (Dilks et al., 2020, 2021; Hahnel et al., 2018). These techniques are difficult in the parasites, and *C. elegans* has been proven to be a suitable model to study resistance complementary to parasitic nematodes (Wit, Dilks, et al., 2021).

New resistance alleles continue to be identified in parasite populations, which emphasizes the need to develop compounds that can be used in conjunction with BZs to potentiate their effects. However, the mechanism of action beyond beta-tubulin binding is still unknown. In susceptible animals, BZs binding to beta-tubulin inhibits tubulin polymerization necessary to form microtubules (Ireland et al., 1979; Lacey, 1990; Lacey & Prichard, 1986; Laclette et al., 1980) but is not well understood in what tissues and specific cells microtubule formation is inhibited. Preliminary studies of BZ susceptibility in *H. contortus* taken from sheep treated with fenbendazole were found to have gross disintegration of the interior intestine, suggesting that BZs target beta-tubulin in the intestine of these animals (Jasmer et al., 2000). We can use genetic tools such as transgenesis and high-throughput assays developed for *C. elegans* to further investigate the mode of action of BZs.

Here, we re-introduced the wild-type *ben-1* gene into a *ben-1* knockout strain background using transgenesis (Rieckher & Tavernarakis, 2017), where multi-copy arrays express *ben-1* in specific tissues. Plasmids containing the coding sequence of *ben-1* fused to tissue-specific promoters, including neurons, hypodermis, muscles, and intestine, as well as endogenous and ubiquitous expression, formed extrachromosomal arrays in transgenic animals. Loss of *ben-1* causes BZ resistance (Hahnel et al., 2018), so transgenic addition of wild-type *ben-1* can restore BZ sensitivity. We then performed high-throughput fitness assays to quantitatively assess the response to albendazole (ABZ). We found that when *ben-1* is expressed in neurons, the wild-type susceptibility phenotype is restored. We then generated transgenic strains that expressed *ben-1* in cholinergic, dopaminergic, GABAergic, or glutamatergic neurons to narrow down the neurons where BZs cause lethality. We found that *ben-1* expression in cholinergic neurons was sufficient to restore wild-type BZ susceptibility. These results offer insights into the mode of action of BZs and suggest that BZs might have a similar cellular target as other classes of anthelmintics.

## 2. Materials and methods

### 2.1 Strains

Strains were maintained at 20°C on modified nematode growth media plates (NGMA) with 1% agar, 0.7% agarose, and *Escherichia coli* OP50 bacteria for food (Andersen et al., 2014). To alleviate starvation effects, strains were grown for three generations before each assay (Andersen et al., 2015). Most strains were generated using the ECA882 strain, *ben-1*(*ean64*), which has the laboratory-derived reference strain background (N2) and a deletion of *ben-1* exons 2 through 4. This strain has been previously shown to be resistant to albendazole (Hahnel et al., 2018). The N2 strain was also used as a background for some control transgenic strains.

### 2.2 Plasmid construction

All *ben-1* plasmids with alternative promoters were constructed by VectorBuilder (Table S1). Tissue-specific promoters included *myo-3* (muscles), *unc-119* (pan-neuronal), *col-19* (hypodermis), and *ges-1* (intestines). A plasmid with the *eft-3* promoter for ubiquitous expression was designed as well. Neurotransmitter-specific promoters included *unc-17* (cholinergic), *eat-4* (glutamatergic), *dat-1* (dopaminergic), and *unc-25* (GABAergic). Promoter sequences for neurotransmitter system-specific expression plasmids have been published previously (Flames & Hobert, 2009; Serrano-Saiz et al., 2020). The plasmid with the *ben-1* endogenous promoter was constructed using the *ben-1* cDNA and an amplicon of the *ben-1* promoter and assembled using Gibson cloning (Gibson et al., 2009). The co-injection marker plasmid, pBCN27 (*myo-2p::GFP::unc-54_3’UTR*) was a gift from Ben Lehner (Addgene, plasmid #26347) (Semple et al., 2010).

### 2.3 C. elegans transgenesis

The *C. elegans* microinjection technique has been previously described (Rieckher & Tavernarakis, 2017). Briefly, a *ben-1* expression plasmid was combined with the *myo-2p::GFP* co-injection marker (5 ng/uL) and 1 kb DNA ladder (Invitrogen #10787018) at the specified concentrations (Table S2, Abrahante et al., 1998; Carvelli et al., 2004; Eastman et al., 1999; Frøkjaer-Jensen et al., 2008; Frøkjær-Jensen et al., 2012; Hao et al., 2012; Kaymak et al., 2016; Marshall & McGhee, 2001; Muñoz-Jiménez et al., 2017; Serrano-Saiz et al., 2013; Thomas et al., 2019). This mixture was injected into the gonads of adult hermaphrodites harboring the *ben-1* deletion (Figure 1). Post-injection, a single adult was placed onto a 6 cm plate. F1 progeny expressing GFP in the pharynx were identified 48-72 hours following injection and subsequently singled (Figure 1). Two independent lines per rescue construct were selected based on the presence of the transgene in the F2 generation (Figure 1). Wild-type GFP control strains were made by injecting the co-injection marker into the N2 strain. Deletion GFP control strains were constructed using the co-injection marker injected into ECA882.

**Figure 1.**
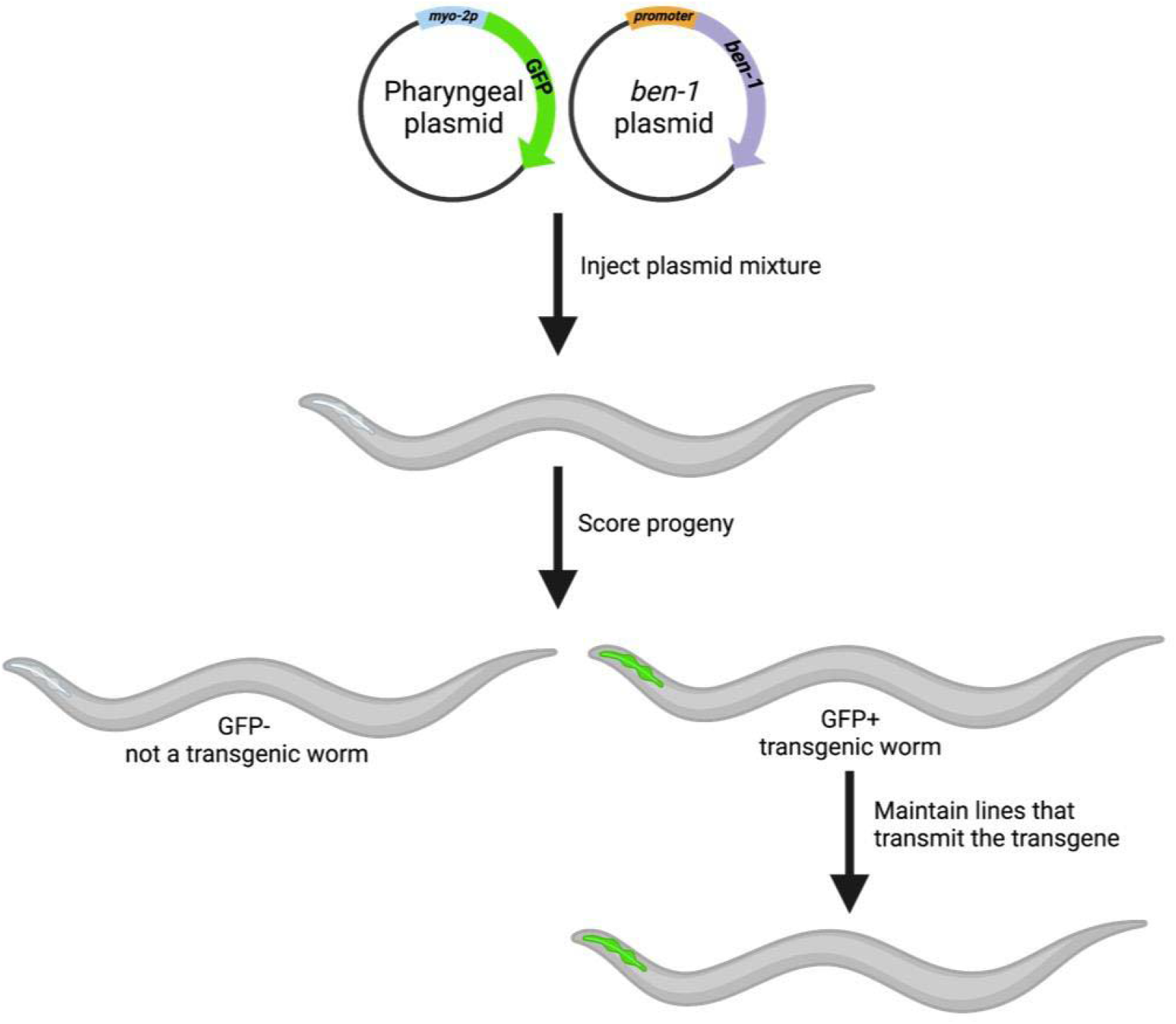
The generation of transgenic strains with specified expression of *ben-1* using microinjection. A mixture of *ben-1* plasmid fused to a specific promoter and a GFP co-injection marker that expressed GFP in the pharynx were injected into the gonads of adult worms. After 48-72 hours, offspring were scored for presence of GFP in the pharynx, which was indicative of successful transgenesis. Strains with approximately 70% transmission were selected for high-throughput fitness assays.

### 2.4 High-throughput fitness assays

The COPAS BIOSORT high-throughput phenotyping assay has been previously described (Andersen et al., 2015; Brady et al., 2019; Dilks et al., 2020, 2021; Evans et al., 2018; Evans & Andersen, 2020; Hahnel et al., 2018; Zdraljevic et al., 2017). Additional measures were taken at each step of the propagation protocol to select for animals with the transgene. Briefly, a small chunk from a starved 6 cm NGMA plate was placed onto a fresh plate. After 48 hours, GFP-positive, gravid hermaphrodites were transferred to a plate with a bleach solution (40 mL NaOCl (Fisher #SS290-1), 10 mL of 10 M NaOH added to 150 mL of distilled water). Approximately 24 hours later, GFP-positive L1 larvae were transferred to a fresh 6 cm NGMA plate. After 48 hours, five animals at the L4 stage were picked to a new plate. After 72 hours to allow for offspring to grow to the L4 stage, five GFP-positive L4s were placed onto a fresh 6 cm NGMA plate and allowed to develop and propagate. After 96 hours, strains were washed off of plates using M9 buffer into 15 mL conicals and treated with fresh bleach solution to dissolve gravid adults and obtain a large number of unhatched embryos. Embryo pools were washed three times using M9 and once using K medium (51 mM NaCl, 32 mM KCl, 3 mM CaCl2, and 3 mM MgSO4 in distilled water) (Boyd et al., 2012) before being resuspended in K medium. Clean embryos were diluted to approximately one embryo per µL in K medium and then aliquoted into 96-well plates at approximately 50 embryos in each well (Figure 2A). After hatching overnight, arrested L1 larvae were fed lyophilized *E. coli* strain HB101 (Pennsylvania State University Shared Fermentation Facility, State College, PA) at a concentration of 5 mg/mL (García-González et al., 2017). After 48 hours, L4 larvae were sorted using the COPAS BIOSORT (Union Biometrica, Holliston MA). The COPAS BIOSORT is able to measure the time-of-flight (TOF) and green fluorescence of each object as it flows through the device (Figure 2B, 2C) (Andersen et al., 2015; Brady et al., 2019; Dilks et al., 2020, 2021; Evans et al., 2018; Evans & Andersen, 2020; Hahnel et al., 2018; Zdraljevic et al., 2017). Three GFP-positive L4 larvae were sorted from each well to wells in a new 96-well plate containing HB101 lysate at 10 mg/mL and 12.5 µM albendazole in 1% DMSO or 1% DMSO alone (Figure 2A). This concentration of albendazole has been previously used with this protocol (Hahnel et al., 2018). Four days after exposure to albendazole, wells were treated with 50 mM sodium azide and scored using the COPAS BIOSORT (Figure 2A).

**Figure 2.**
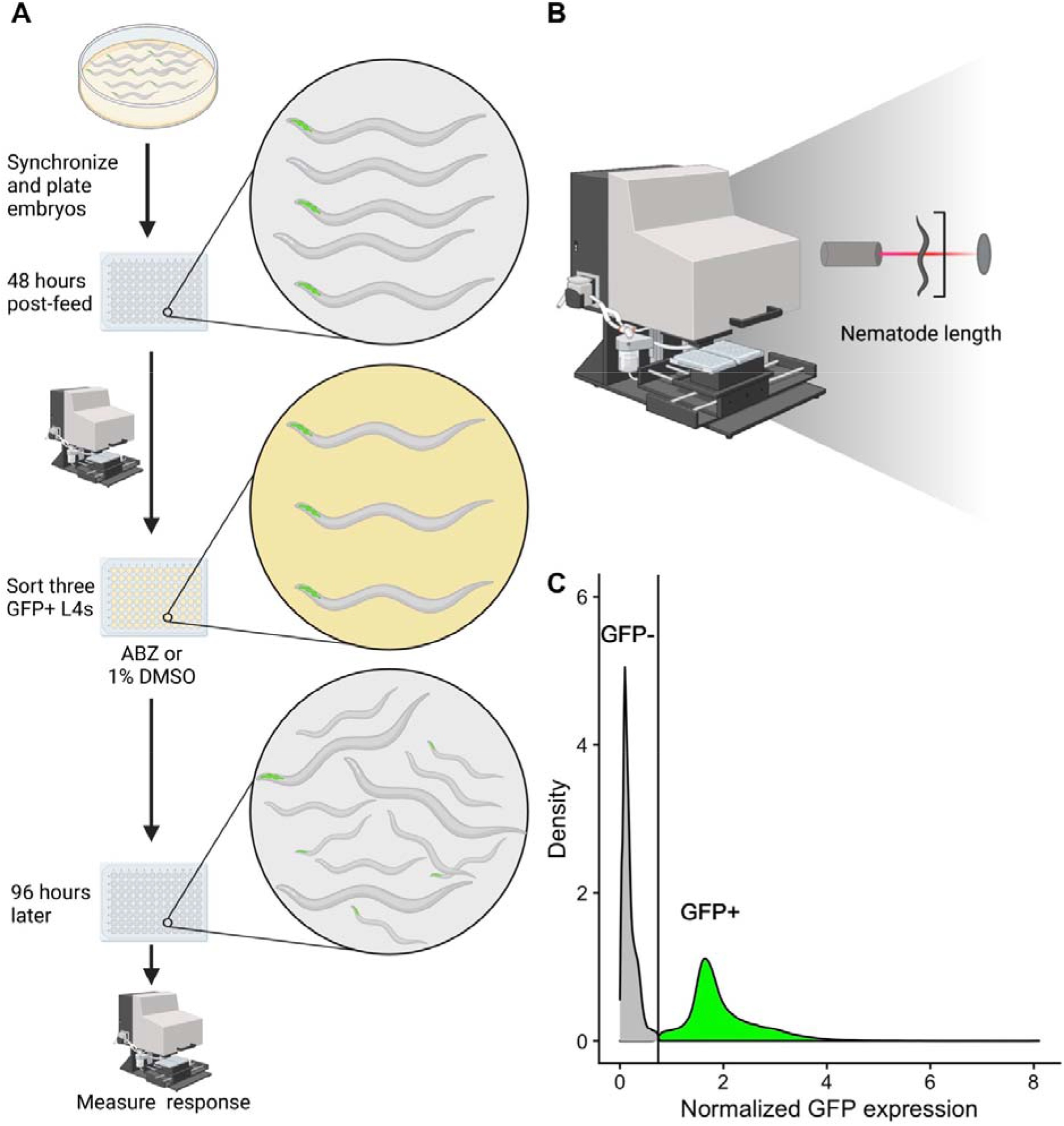
COPAS BIOSORT high-throughput fitness assay. **A)** Illustration of the high-throughput fitness assay is shown. Synchronized worms were treated with bleach solution and embryos were distributed into 96-well plates. 48 hours post-feeding, three GFP-positive L4 animals were sorted using the COPAS BIOSORT into each well of a 96-well plate containing 12.5 µM albendazole in 1% DMSO or 1% DMSO alone. After 96 hours, animals were scored for GFP fluorescence and length using the COPAS BIOSORT. **B)** Illustration of COPAS BIOSORT is shown. Animals were passed through a flow cell and a laser. The time the detector is interrupted is equated to the length of the nematode. Illustration adapted from (Wit, Rodriguez, et al., 2021). **C)** Distribution of GFP fluorescence of animals is shown. The x-axis is the GFP expression of each animal normalized by the length of each animal. The y-axis is the distribution of the population. The vertical line represents the threshold for establishing if animals are GFP-positive (green) or GFP-negative (gray).

### 2.4 Data analysis

Raw data from the COPAS BIOSORT were processed using the R package *easysorter* (Shimko & Andersen, 2014) as previously described (Dilks et al., 2020, 2021; Hahnel et al., 2018). An average value for each phenotypic trait measured in the control condition (1% DMSO) was deducted from the albendazole condition data to normalize the data for each strain. The distribution of green fluorescence values was analyzed to establish a threshold to filter for GFP-positive animals that had the transgene (Figure 2C). The average mean TOF value was summarized for each well per strain and the distribution of mean TOF values for each transgenic strain was compared to the mean TOF values for the *ben-1* deletion strain. Statistical tests and analyses were performed in R using the *tukeyHSD* function in the *Rstatix* package. The ANOVA model (*phenotype ∼ strain*) was used to compare differences in phenotypic responses to BZs between the *ben-1* deletion strain and the other strains.

### 2.5 Data availability

A list of plasmids used with vendor ID information and a list of *C. elegans* strains and genotypes used in experiments are included as supplementary information (Tables S1 and S2). The data and code used to process these data are available at https://github.com/AndersenLab/2022_ben1sensitivity_SBG.

## 3. Results

### 3.1 ben-1 function in neurons rescues BZ sensitive phenotype

We created transgenic strains where *ben-1* was expressed in different tissues, including muscles, neurons, hypodermis, intestines, as well as ubiquitous expression. These different expression constructs were made in the ECA882 *ben-1* deletion background. For each candidate tissue-specific expression strain, two independent strains were generated and assayed to ensure that the measured effects were caused by the transgenes and not a vagary of the injection process. Assay results with the second strain are available as supplemental information (Figures S1 and S2). Animals with a *ben-1* specific transgene were identifiable by a green, fluorescent pharynx caused by the pharyngeal expression of the co-injection transgenesis marker. We then performed high-throughput assays to test if any of the tissue-specific expression strains rescued susceptibility to albendazole. A strain with the resistant *ben-1* knockout background and the pharyngeal expression marker and a strain with the susceptible wild-type background and the pharyngeal expression marker were used as controls. We measured growth in control conditions (DMSO) and albendazole (ABZ) conditions in 44 replicates per strain. The high-throughput assay was performed as described (See 2.4, figure 2A). Following 48 hours of growth, three GFP-positive animals for each strain were sorted into each well of a 96-well plate using the COPAS BIOSORT. These animals gave rise to a population in each well that was then scored 96 hours after exposure to ABZ using the COPAS BIOSORT. The green fluorescence measurements were used to filter the data to only include animals that had the transgene (Figure 2C). We used the length of animals as a measure of developmental rate. Strains with lower average length values, when compared to the *ben-1* deletion strain, indicate susceptibility to ABZ. As expected, we measured a significant difference in animal length between the wild-type strain as compared to the *ben-1* deletion in response to ABZ. (Figure 3A). We found that expression of *ben-1* under its endogenous promoter or when highly expressed in all tissues caused susceptibility to ABZ (Figures 3B and S1). Furthermore, the high level of expression using the *eft-3* promoter caused ABZ susceptibility far beyond wild-type levels. When *ben-1* expression was driven solely in neurons using the *unc-119* promoter, the animals were equally susceptible to ABZ as the strains that had endogenous expression of *ben-1*. These results suggest that neuronal *ben-1* might be the endogenous target of BZ compounds. Expression of *ben-1* in hypodermis, muscles, and intestine did not restore the ABZ susceptibility phenotype (Figure 3B), indicating that neurons might be the sole target of BZs.

**Figure 3.**
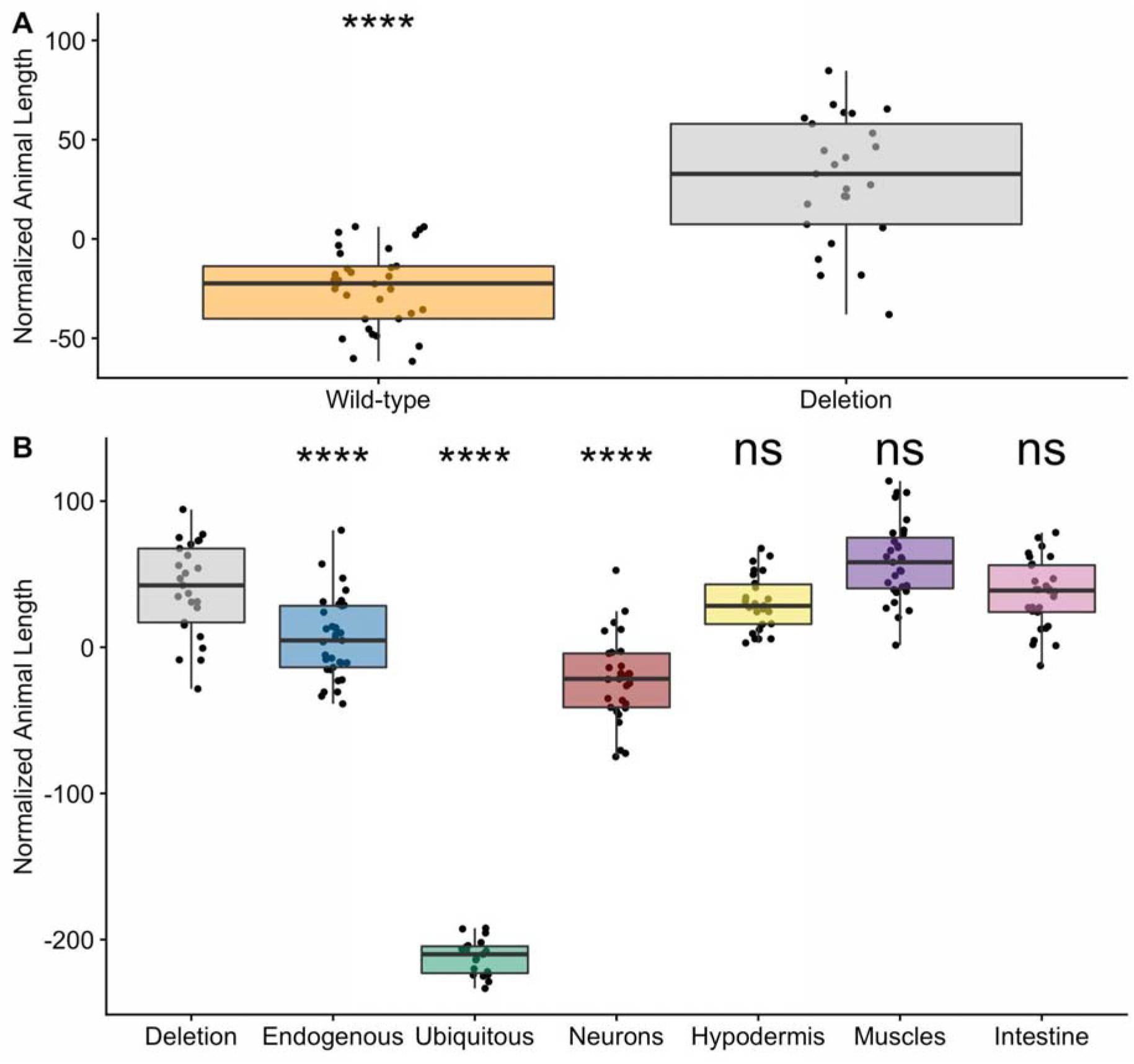
*ben-1* function in neurons underlies BZ sensitivity. **A)** The x-axis denotes the genetic background at the *ben-1* locus for each strain. The y-axis is the normalized length of animals after exposure to 12.5 µM of albendazole. Phenotypic data was normalized by subtracting an average value for each trait from the control data. Each data point is the mean value for the population of GFP-positive worms in a single well. Tukey box plots have horizontal lines at the third quartile on the top, the median in the middle and the first quartile at the bottom. The whiskers are extended within 1.5 range from each quartile. The significant difference between the wild-type genotype and the deletion genotype is shown above the wild-type results (**** = p < 0.0001, one-way ANOVA, Tukey HSD). **B)** The x-axis denotes the specificity of the *ben-1* expression from the transgene. The y-axis is the normalized length of animals after exposure to 12.5 µM of albendazole. Phenotypic data was normalized by subtracting an average value for each trait from the control data. Each data point is the mean value for the population of GFP-positive worms in a single well. Tukey box plots have horizontal lines at the third quartile on the top, the median in the middle and the first quartile at the bottom. The whiskers are extended within 1.5 range from each quartile. The significant difference between the transgene and the deletion genotype is shown above the transgene (**** = p < 0.0001, one-way ANOVA, Tukey HSD)

### 3.2 ben-1 function in cholinergic and GABAergic neurons rescues BZ sensitive phenotype

Although beta-tubulin genes are expressed in every cell of an organism, *ben-1* has been specifically shown to be broadly expressed in neurons (Hurd, 2018). We used the *C. elegans* Neuronal Gene Expression Map and Network (CeNGEN) dataset (Taylor et al., 2021) to further explore neuron-specific expression of *ben-1* and found that it is expressed in 97 of the 128 cell types distinguished in the dataset and primarily in cholinergic and glutamatergic neurons (Figure S3). The thorough characterization of each neuron in the *C. elegans* nervous system offered the opportunity to generate transgenic animals with *ben-1* expression specific to subsets of neurons. We hoped to narrow down which neurons are specifically targeted by BZs by creating transgenic strains with *ben-1* expression separated by neurotransmitter system and perform the same high-throughput analyses.

We created transgenic strains where *ben-1* expression in neurons was subsetted by the neurotransmitter system, including cholinergic, glutamatergic, dopaminergic, and GABAergic neurons. Two independent lines for each type were generated in the ECA882 *ben-1* deletion background. The same pharyngeal expression marker was used to identify transgenic animals as well as the same wild-type and resistant control strains. We compared the responses of the four neurotransmitter system-specific transgenes as well as the endogenous and pan-neuronal expression strains to the two control strains with 44 replicates per strain. Offspring were scored 96 hours after three GFP-positive parents were sorted into each well of plates containing either ABZ or the control DMSO condition. The score data were filtered to only compare the lengths of GFP-positive animals. We found that animals expressing *ben-1* in cholinergic regions were significantly smaller (*e.g*., developmentally delayed because of susceptibility to ABZ) than the resistant strain when exposed to ABZ and were comparable in size to animals expressing *ben-1* in all neurons (Figure 4). It is worth noting that a large proportion of the neuron classes that express *ben-1* are cholinergic, including a few of the neurons with the highest levels of expression (Figure S3, Loer & Rand, 2022, Taylor et al., 2021). We observed no difference in animal length between the resistant strain and animals expressing *ben-1* in glutamatergic and dopaminergic neurons, suggesting that BZs do not target these neurons (Figure 4). In animals that expressed *ben-1* in GABAergic neurons, we found a less extreme ABZ response (Figure 4), suggesting that *ben-1* function in cholinergic and potentially GABAergic neurons was sufficient to restore ABZ susceptibility.

**Figure 4.**
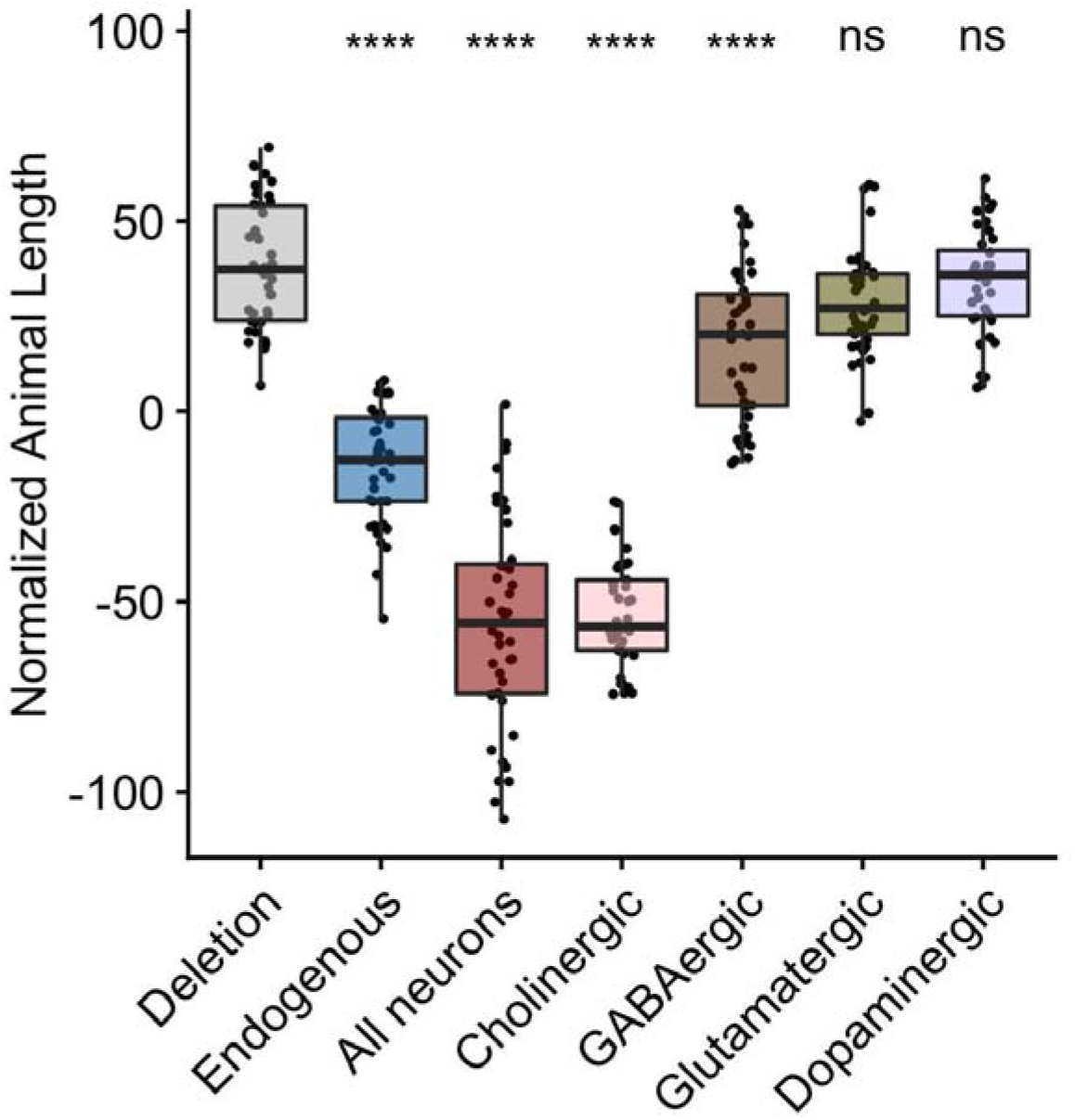
*ben-1* function in cholinergic and GABAergic neurons underlies BZ sensitivity. The x-axis denotes the specificity of the *ben-1* expression from the transgene. The y-axis is the normalized lengths of animals after exposure to 12.5 µM of albendazole. Phenotypic data was normalized by subtracting an average value for each trait from the control data. Each data point is the mean value for the population of GFP-positive worms in a single well. Tukey box plots have horizontal lines at the third quartile on the top, the median in the middle and the first quartile at the bottom. The whiskers are extended within 1.5 range from each quartile. The significant difference between the transgene and the deletion genotype is shown above the transgene (**** = p < 0.0001, one-way ANOVA, Tukey HSD)

## 4. Discussion

### 4.1 ben-1 function in cholinergic and GABAergic neurons is sufficient to rescue organismal BZ susceptibility

Although it is well understood that reduction- or loss-of-function variation in *ben-1* confers resistance to BZs (Driscoll et al., 1989; Hahnel et al., 2018), little has been done to better characterize BZ sensitivity at the cellular level. Understanding cell-type specific targets of BZs could enable development of co-treatments that potentiate BZ effects. Here, we generated transgenic *C. elegans* where *ben-1* expression was limited to one of four tissue types or ubiquitous expression and compared to endogenous expression. Each strain’s response to the BZ drug albendazole (ABZ) was compared to ABZ responses in the resistant *ben-1* deletion strain. When *ben-1* is expressed only in neurons, we see that animal development is delayed in ABZ conditions (Figure 3), suggesting that *ben-1* expression in neurons is sufficient to rescue sensitivity to BZs. Although in the replicate experiment we also found a statistically significant difference in animal length when *ben-1* was expressed in the hypodermis or the intestine (Figure S1), these results were not reproducible between experiments so could be attributed to the overexpression of genes by different extrachromosomal arrays. These experiments all rely on the overexpression of *ben-1* in a tissue-specific manner. This overexpression could cause an imbalance in endogenous levels of beta-tubulins and could cause a slight deleterious effect on fitness. The normalized length values for strains expressing *ben-1* in the hypodermis and the intestine are also close to zero or positive, unlike the negative normalized values for animals expressing *ben-1* in neurons where the difference between the response of the resistant strain is much more pronounced (Figures 3B, S1). Single-cell expression studies show that *ben-1* is primarily expressed in neurons (Hurd, 2018), so neurons might require this specific form of beta-tubulin. Future experiments can confirm if *ben-1* function in neurons is necessary for the BZ susceptibility phenotype.

The detailed understanding of the *C. elegans* nervous system anatomy and function allowed us to identify specific neurons that are targeted by BZs (Taylor et al., 2021; White et al., 1986). We subsetted the 118 neuron classes by neurotransmitter system to determine which neurons might require *ben-1* function to be susceptible to BZ compounds. Transgenic strains were generated with *ben-1* fused to cholinergic, dopaminergic, GABAergic, or glutamatergic-specific promoters (Flames & Hobert, 2009; Serrano-Saiz et al., 2020) and assayed like the tissue-specificity experiments. When *ben-1* is expressed in cholinergic neurons, the drug-treated animals were significantly smaller than the resistant strain, similar to when expression is driven in all neurons (Figure 4). Although some glutamatergic and dopaminergic neurons also have higher levels of *ben-1* expression, a significant phenotypic effect was not observed in animals driving *ben-1* expression in these neurons (Figures 4 and S2). However, it is possible that BZs also target GABAergic neurons as a less-significant deleterious effect was consistently observed across experiments (Figures 4 and S2). Neuron expression data of *ben-1* supports that cholinergic neurons are more likely to be the target of BZs because they express *ben-1* at higher levels than the few GABAergic neurons with *ben-1* expression (Loer & Rand, 2022, Taylor et al., 2021).

### 4.2 Further investigation of ben-1 function in specific tissues is required

*C. elegans* transgenesis using microinjection of plasmids containing *ben-1* is an effective model to suggest where *ben-1* function might cause susceptibility to BZs. The re-introduction of *ben-1* as an extrachromosomal array causes overexpression, which might not reflect endogenous levels and can cause artifactual conclusions. To definitively show the role of *ben-1* in specific neurons, we require experiments to test the requirement of *ben-1* in BZ resistance. Conditional knockout (cKO) experiments where the *ben-1* gene is removed from wild-type animals in specific tissues and/or cells is achievable using the *Cre/loxP* system developed for *C. elegans* (Kage-Nakadai et al., 2014). Genes flanked with *loxP* sites are removed from the genome using the *Cre* recombinase (Austin et al., 1981; Kage-Nakadai et al., 2014). Conditionally expressed *Cre* will only excise *ben-1* from specific target tissues and/or cells, and the remaining tissues of the organism will express *ben-1* endogenously. We can create strains with *ben-1* knocked out in specific tissue types or neurons and perform the same high-throughput assays to assess the relative fitness of each strain when exposed to BZs. If animals with *ben-1* knocked out only in neurons are resistant to BZs, *ben-1* function in this tissue is necessary for BZ sensitivity. Neuron-specific knockout experiments will confirm specific targets of BZs.

### 4.3 Identify which specific neurons are targeted by BZs

With the suggestion of *ben-1* function in cholinergic and GABA neurons underlying BZ sensitivity, we can further narrow down the neurons targeted by BZs. We can continue to generate transgenic worms using promoters for genes specific to different neuron classes and leverage other tools unique to *C. elegans* such as the NeuroPAL strain set (Yemini et al., 2021) and the CRF_ID annotation framework (Chaudhary et al., 2021). NeuroPAL (Neuronal Polychromic Atlas of Landmarks) strains are transgenic animals with a differentiated fluorescence pattern for every neuron in the organism (Yemini et al., 2021). Specialized software was developed for the system that can be used to identify each neuron from images based on fluorescence (Yemini et al., 2021). However, the annotation software is semi-automatic and requires some manual input to label microscopy images (Yemini et al., 2021). An automated annotation framework called CRF_ID has been developed to label cells with an algorithm based on the graphical-model based framework Conditional Random Fields (CRF) (Chaudhary et al., 2021). It has been shown to improve accuracy in labeling whole-brain images from *C. elegans* and be compatible with labeling color-based NeuroPAL animal images (Chaudhary et al., 2021). This sophisticated imaging and annotation system would allow us to simultaneously assay the entire *C. elegans* nervous system to identify specific targets of BZs. Neurons affected by BZs would be identified by comparing NeuroPAL animals treated with BZs to ones developing in control conditions. The ability to assay the entire nervous system will be particularly useful as not all of the 118 neuron classes were accounted for in the neurotransmitter system-specific assays as a few neurons expressing *ben-1* use other neurotransmitters or their neurotransmitter system is unknown (Figure S3).

### 4.4 Benzimidazole neuron targets might be shared with other anthelmintics

The finding of BZs targeting beta-tubulin in neurons suggests that the specific neurons targeted might be in common with other anthelmintic compounds. Resistance to two other widely used drug classes, macrocyclic lactones (MLs) and nicotinic acetylcholine receptor agonists (nAChRs), has been associated with genes expressed in neurons. Resistance to MLs has been linked to genes coding for the subunits of glutamate-gated chloride channels, including *glc-1, avr-14*, and *avr-15* (Dent et al., 2000), and genes involved in resistance to nAChRs, including *unc-63, unc-38, unc-29, lev-1*, and *lev-8* are known (Lewis et al., 1980; Qian et al., 2008). Analyzing overlaps in expression between ML and nAChR resistance genes and *ben-1* using the CeNGEN data set (Taylor et al., 2021) could identify potential neurons targeted by multiple anthelmintic compounds. Comparing responses NeuroPAL animals have to each compound could confirm if some neurons might be targeted and inhibited/killed by multiple anthelmintic drugs. Identification of neurons that are targeted by multiple anthelmintics might allow for the development of co-treatments compatible with more than one anthelmintic compound. Overall, the development of sophisticated analytical systems for assaying the entire *C. elegans* nervous system has the potential to improve our understanding of mechanisms of action for multiple anthelmintic classes.

## Acknowledgements

We would like to thank Clay Dilks, Katie Evans, and Nicole Banks for their help with experiments and analysis and Amanda Shaver for her helpful comments on this manuscript. This work was supported by the National Institutes of Health NIAID grant R01AI153088 to ECA. This study used data made available by the *C. elegans* Neuronal Gene Expression Map and Network (NIH NINDS R01NS100547). Neurotransmitter system data are from WormAtlas neurotransmitter tables by Curtis M. Loer and James B. Rand (Loer & Rand, 2022, compiled from Gendrel et al., 2016; Pereira et al., 2015; Serrano-Saiz et al., 2017). Some diagrams in figures were created using BioRender.com.

## Supplementary figures and legends

**Figure S1.**
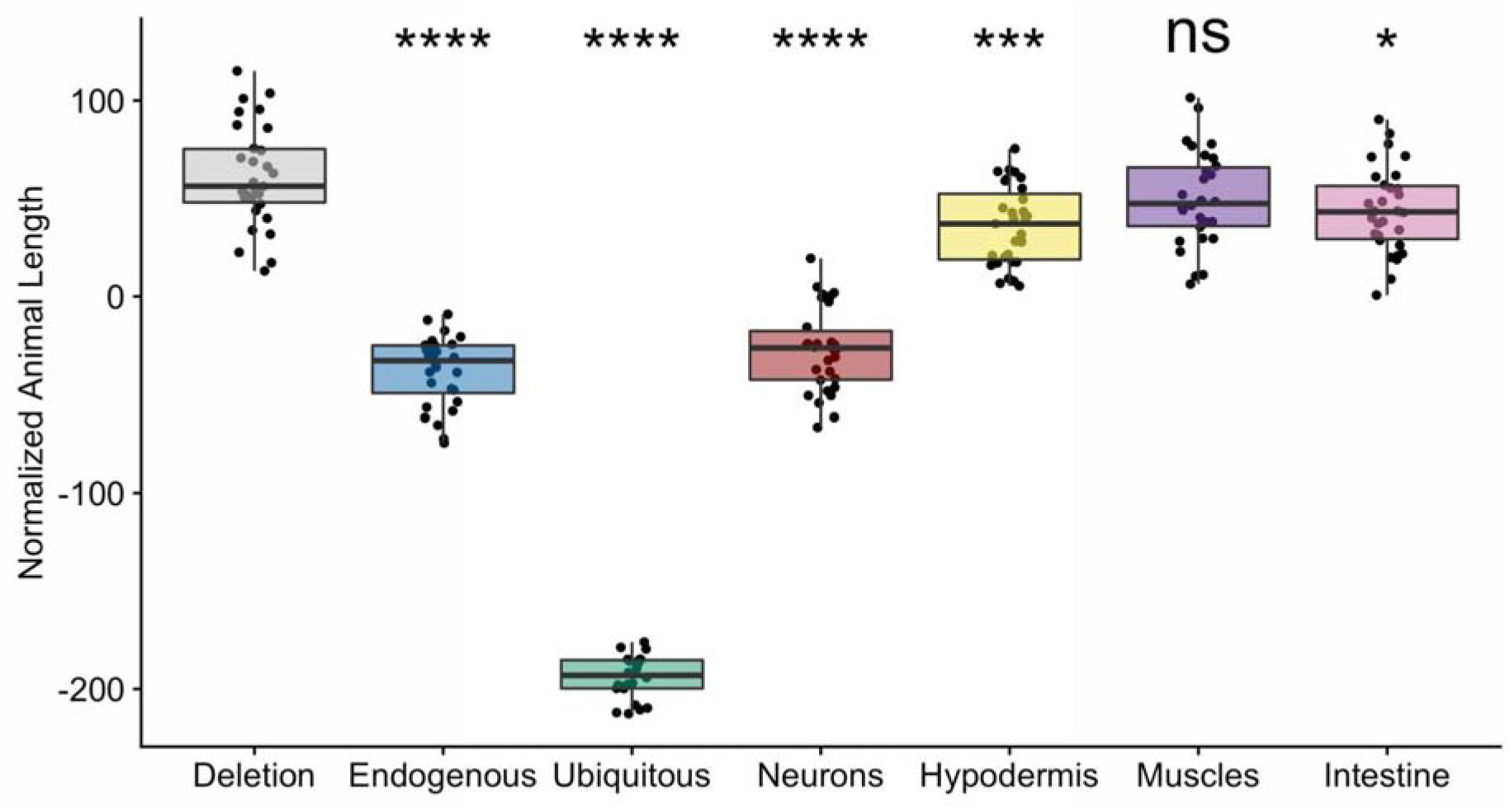
Second independent line results. The x-axis denotes the specificity of the *ben-1* expression from the transgene. The y-axis is the normalized lengths of animals after exposure to 12.5 µM of albendazole. Phenotypic data was normalized by subtracting an average value for each trait from the control data. Each data point is the mean value for the population of GFP-positive worms in a single well. Tukey box plots have horizontal lines at the third quartile on the top, the median in the middle, and the first quartile at the bottom. The whiskers are extended within 1.5 range from each quartile. The significant difference between the wild-type genotype and the deletion genotype is shown above the wild-type results (* =p < 0.05, *** = p < 0.001, **** = p < 0.0001, one-way ANOVA, Tukey HSD).

**Figure S2.**
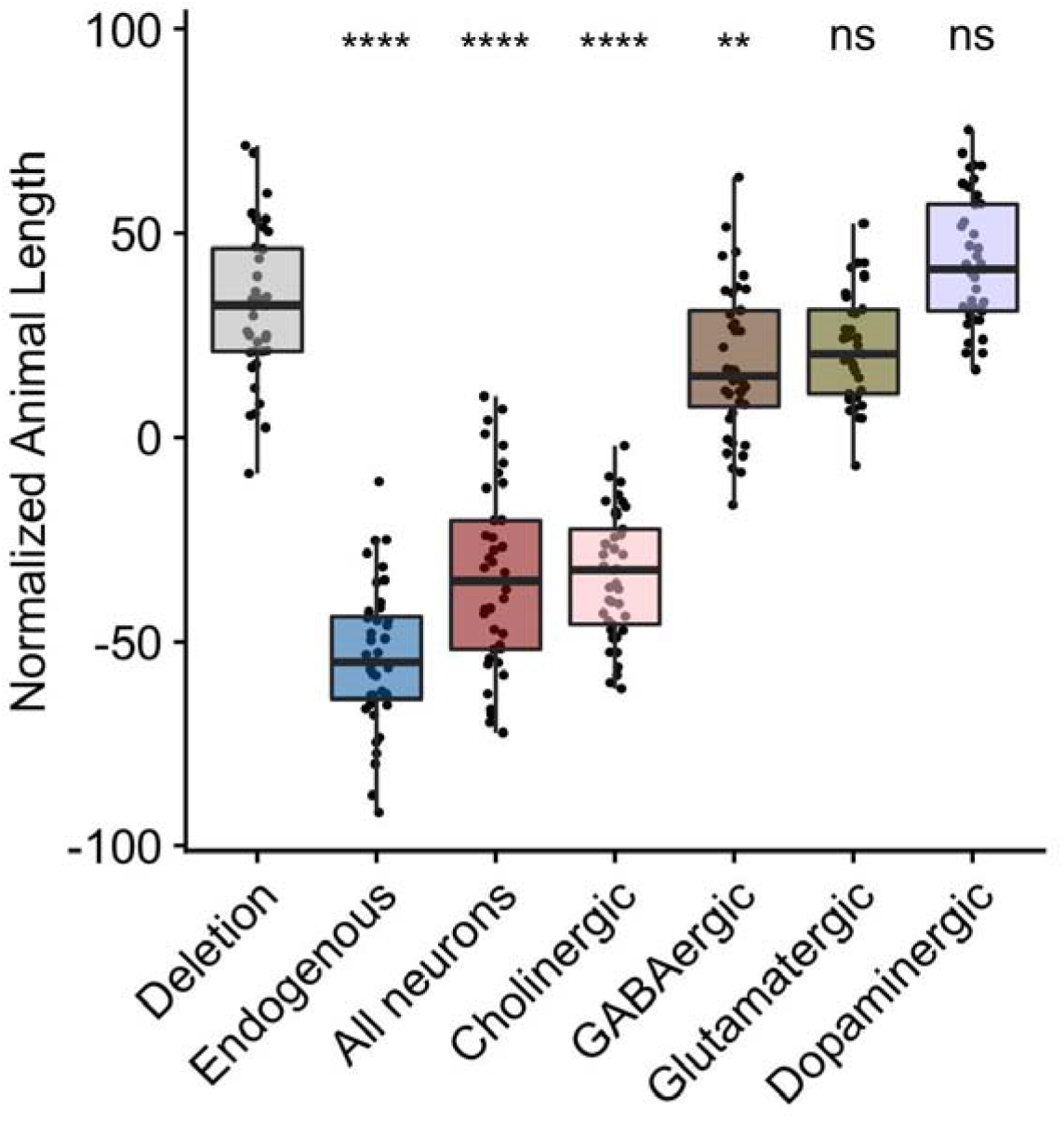
Second independent line results. The x-axis denotes the specificity of the *ben-1* expression from the transgene. The y-axis is the normalized lengths of animals after exposure to 12.5 µM of albendazole. Phenotypic data was normalized by subtracting an average value for each trait from the control data. Each data point is the mean value for the population of GFP-positive worms in a single well. Tukey box plots have horizontal lines at the third quartile on the top, the median in the middle, and the first quartile at the bottom. The whiskers are extended within 1.5 range from each quartile. The significant difference between the wild-type genotype and the deletion genotype is shown above the wild-type results (** = p < 0.01, < **** = p < 0.0001, one-way ANOVA, Tukey HSD).

**Figure S3.**
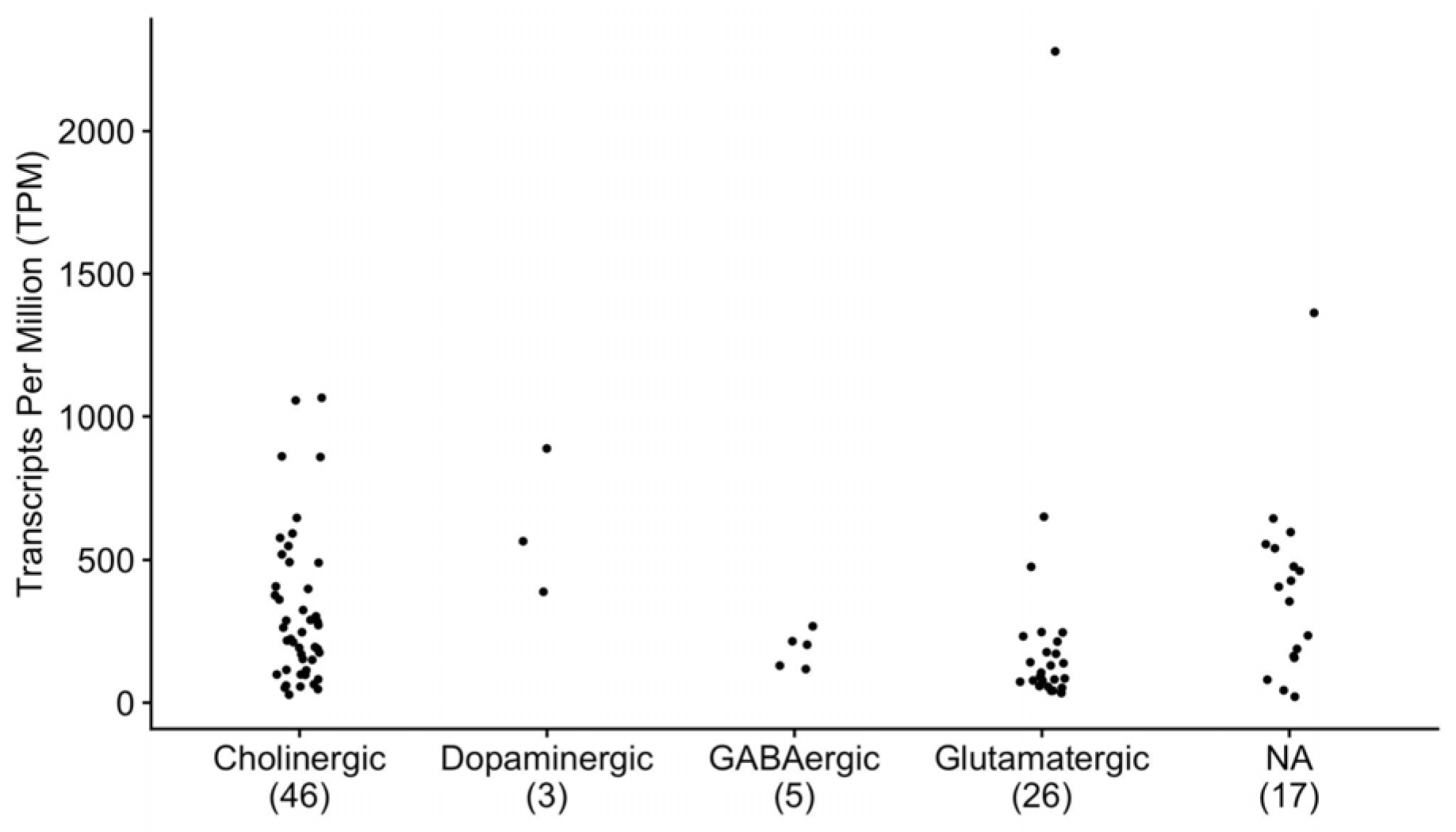
Distribution of neurotransmitter systems for neuron types expressing *ben-1*. The expression data used for this figure are from the CeNGEN data set (Taylor et al., 2021). Neurotransmitter type data are from WormAtlas neurotransmitter tables by Curtis M. Loer and James B. Rand (Loer & Rand, 2022, compiled from Gendrel et al., 2016; Pereira et al., 2015; Serrano-Saiz et al., 2017). The x-axis denotes the neurotransmitter system for each neuron. The number of neuron classes in each group is included below the neurotransmitter system. NA denotes neurons where the neurotransmitter is unknown or not one of four neurotransmitter systems was included in our experiments. The y-axis is the expression level of *ben-1* for each neurotransmitter type measured as transcripts per million (TPM).

